# Urinary Metabolomics from a Dose-Fractionated Polymyxin B Rat Model of Acute Kidney Injury

**DOI:** 10.1101/2021.05.21.444980

**Authors:** Emanuela Locci, Jiajun Liu, Gwendolyn M. Pais, Alberto Chighine, Dariusc Andrea Kahnamoei, Theodoros Xanthos, Athanasios Chalkias, Andrew Lee, Alan R. Hauser, Jack Chang, Nathaniel J. Rhodes, Ernesto d’Aloja, Marc H. Scheetz

**Author notes:** Corresponding Author: Emanuela Locci, Università degli studi di Cagliari | UNICA · Department of Medical Sciences and Public Health. at the time the work was conducted (Jiajun Liu).

## Abstract

**Background:** Polymyxin B remains an important antimicrobial against multi-drug resistant bacteria; however, kidney injury is often a treatment limiting event with kidney failure rates that range from 5-13%.

**Methods:** Samples were obtained from a previously conducted study of male Sprague-Dawley rats that received dose fractionated polymyxin B (12 mg/kg/day subcutaneously) once daily (QD), twice daily (BID), and thrice daily (TID) for three days. In the original study, urinary biomarkers and kidney histopathology scores were determined. Urine was sampled daily and analyzed for urinary metabolites via 1H NMR analysis.

Unsupervised Principal Components Analysis was applied for exploratory data analysis to identify trends and outliers in the spectral data. Then, supervised Orthogonal Partial Least Square Discriminant Analysis was applied to classify the samples collected in different days and identify metabolic differences during the treatment. Metabolomes were compared across study groups (i.e. those receiving QD, BID, TID, and control) using a mixed-effects models. Spearman correlation was performed for injury biomarkers and the metabolome.

**Results:** A total of 27 rats contributed 77 urinary samples; n=25 rats were included that were treated with Polymyxin B and n=2 received saline. Pre-dosing samples clustered well and were characterized by higher amounts of citrate, 2-oxoglutarate, and Hippurate. On day 1 post treatment, day 1 samples showed higher taurine; day 3 samples had higher lactate, acetate and creatine. Taurine was the only metabolite significantly increased in both BID and TID compared to QD group. Taurine on day 1 correlated with increasing histopathology scores (Spearman’s rho = 0.4167, P=0.038) and KIM-1 (Spearman’s rho =0.4052, P=0.036); whereas KIM-1 on day one and day 3 did not reach significance with histopathology (Spearman’s rho = 0.3248, P=0.11 and Spearman’s rho = 0.3739, P=0.066).

**Conclusion:** Polymyxin B causes increased amounts of urinary taurine on day 1 which then normalizes to baseline concentrations. Taurine may provide one of the earlier signals of acute kidney damage caused by polymyxin B.

## Background

While antibiotic drug development has increased slightly in recent years, it is projected that novel therapeutics will not keep pace with the rate of antimicrobial resistance.^1^ Without new effective and safe therapeutics, polymyxin agents will remain last line therapeutics for resistant Gram-negative infections and retain a significant portion of use worldwide because of their retained antibiotic activity, world-wide availability, and cost.^2^ Of the systemically utilized polymyxins, polymyxin B (PB) has the more predictable drug exposure and outcome profile (both for efficacy and toxicity); however, kidney injury is very common with PB use, and kidney injury is often a treatment limiting event with kidney failure rates that range from 5-13%.^3^ Although this is well-recognized, the mechanism and pathophysiological pathway of polymyxin-associated nephrotoxicity is still being discerned. After PB is filtered by the glomeruli, it preferentially accumulates in renal proximal tubule cells^3^ via receptor-mediated endocytosis by megalin and possibly by PEPT2.^4–6^ Quantitative mapping of PB demonstrates that accumulation in the kidney is both concentration- and time-dependent (as shown by FADDI-096 uptake).^4^ As with other carrier-mediated transport mechanisms, accumulation seems to be saturable.^7^ Ultimately, accumulation of PB in renal tubule cells causes activation of apoptotic pathways such as caspase activation, endoplasmic reticulum stress, mitochondrial damage, oxidative insult, and cell cycle arrest.^8, 9^ Acute tubular damage then leads to functional decline in creatinine clearance (CrCL), as well as increased serum urea and creatinine concentrations.^10^

The toxicity of PB has been related to plasma 24-hour area under the curve (AUC_24h_), which is a function of both concentration and time.^11^ Studies that have fractionated total daily dosage schemes (to maintained isometric AUCs) have resulted in mixed toxicodynamic outcomes.^12, 13, 14, 15^ In humans, fractionated dosing schemes demonstrated slightly less nephrotoxicity than once daily dosing schemes.^6^ In rats, once daily dosing was associated with less nephrotoxicity as assessed by serum creatinine (SCr), fewer histopathologic lesions, and lower drug concentrations in kidney tissue.^12^ Others conducting similar experiments came to similar conclusions, i.e. rats receiving fractionated dosing schemes had slightly higher degrees of kidney injury than those receiving once daily administration.^14, 15^ AKI was very common in all of the human and preclinical studies, and kidney injury rates and extents were reasonably similar between groups. Thus, it is unlikely that toxicity will be ameliorated by simply changing the dosing scheme (of the same total daily dose); attention must be focused on early identification and prevention of toxicity.

Several pathways have been investigated to reduce polymyxin-associated nephrotoxicity; these include decreasing the uptake of PB by renal tubular cells, attenuating polymyxin-induced oxidative stress, modifying the polymyxin structure, and earlier identification of associated nephrotoxicity. Ameliorating agents for polymyxin-associated nephrotoxicity have been investigated in animal models and clinical trials, including N-acetylcysteine, ascorbic acid, melatonin, and methionine. ^16–19^ However, results have been underwhelming to date. Attempts at detecting polymyxin-associated nephrotoxicity have largely been limited by conventional biomarkers. Serum creatinine is the most commonly used clinical biomarker for identification and staging of AKI; however, serum creatinine is a non-specific marker that is associated with significant delay in the identification of AKI.^9^ After an initial renal insult, function can be reduced by 30-40% for up to 72 hours, prior to a detectable rise in serum creatinine.^20, 21^ Furthermore, even small short-term increases of serum creatinine (e.g. 0.3 mg/dL) have been associated with greater than 3-fold increases in 30-day mortality.^22^ Therefore, new strategies to identify AKI early are necessary to prevent downstream toxicity when patients are treated with therapies such as PB. Novel urinary biomarkers of renal injury such as kidney injury molecule-1 (KIM-1), NGAL, and osteopontin have shown promise in animal models for early detection of AKI.^34^

The field of metabolomics also holds significant potential for precision medicine and earlier identification of disease states. Recent studies have investigated the metabolomics related to AKI in the hopes of earlier and more specific identification.^13^ These studies were similar in design and consist of exploratory efforts to characterize urinary metabolomic biomarkers related to chronic renal disease and specific nephrotoxins such as cisplatin and gentamicin.^35,36,37^ Yet these approaches are still new and the urinary metabolomic profile of polymyxins remain poorly characterized. Early identification of AKI in patients receiving polymyxin could allow clinicians to stop or change therapy before irreversible toxicity results. The purpose of this study was to identify metabolomic markers and signatures associated with PB kidney injury by analyzing samples from a previously conducted rat toxicology study.^14^

## METHODS

Samples were obtained from a previously conducted study and were analyzed in batch as described below.^14^ In brief, male Sprague-Dawley rats (n=32) underwent double jugular catheterization surgery and then received dose fractionated PB at a dose of 12 mg/kg/day subcutaneously once daily (QD), twice daily (BID), and thrice daily (TID) for three days (i.e. day 1, day 2, and day 3). Control groups animals received normal saline (NS). Rats were placed in metabolic cages for the duration of the experiment (i.e. pre-drug/pre-surgery and daily after the drug dosing protocol was initiated for a period of 3 days). Urine was sampled daily and analyzed for a variety of kidney injury biomarkers including kidney injury molecule-1 (KIM-1) which was the most predictive urinary biomarker from the previous study. ^14^ Histopathology was scored according to standard criteria.^23^ The collected rat urine samples were stored at −80°C (except during batch shipment on dry ice) until analysis. In this sub-study, NMR experiments were performed on: 25 PB treated rat urines collected pre-surgery (day 0), post 1^st^ day of dosing (day 1) and post 3^rd^ day of dosing (day 3) and 2 NS treated rat urines collected at day 1 only. The study was reviewed and approved by the Midwestern University Institutional Animal Care and Use Committee.

For ^1^H NMR analysis, thawed urines were centrifuged at 16,750 g for 10 minutes at 4°C and supernatants were added with a 0.1% w/w aqueous solution of sodium azide (NaN_3_) to avoid bacterial growth. 630 μl of each sample were mixed with 70ul of a 1.5 M phosphate buffer solution (pH=7.4) in D_2_O (99.9%, Cambridge Isotope Laboratories Inc, Andover, USA) containing the internal standard sodium 3-(trimethylsilyl)propionate-2,2,3,3,-*d*_*4*_ (TSP, 98 atom % D, Sigma-Aldrich, Milan) at a 0.6 mM final concentration. 650 μl of the obtained solutions were transferred into 5 mm NMR tubes.

^1^H NMR experiments were carried out on a Varian UNITY INOVA 500 spectrometer (Agilent Technologies, CA, USA). ^1^H NMR spectra were recorded using a 1D-NOESY pulse sequence with a mixing time of 1 ms and a recycle time of 3.5 s, for water suppression. Spectra were acquired at 300K, with a spectral width of 6000 Hz, a 90° pulse, and 128 scans. Spectra were processed using MestReNova software (Version 14.1.2, Mestrelab Research S.L.). Before Fourier transformation, the free induction decays (FID) were zero-filled to 128k and multiplied by an exponential weighting function corresponding to a line-broadening factor of 0.5 Hz. Spectra were phased and baseline corrected. Chemical shifts were referred to the TSP single resonance at 0.00 ppm. Spectra were reduced to consecutive integrated spectral regions (bins) of 0.01 ppm width in the region 0.80 - 9.50 ppm. The spectral region between 4.70 and 4.98 ppm containing the residual water resonance was excluded from the analysis. To minimize the effects of variable concentration among different samples, the integrated area within each bin was normalized to a constant sum of 100 for each spectrum. The assignment of major resonances was performed on the basis of data published in the literature and using the database implemented in Chenomx NMR Suite 8.2 Library (Chenomx Inc., Edmonton, Canada).^24^ The final data matrix was imported into the SIMCA software (Version 14.0, Umetrics, Umeå, Sweden) for multivariate statistical analysis. Pareto scaling was applied prior to perform the analysis.

Unsupervised Principal Components Analysis (PCA) was applied for exploratory data analysis to identify trends and outliers in the spectral data. Then, supervised Orthogonal Partial Least Square Discriminant Analysis (OPLS-DA) and its variant (OPLS-DA) was applied to classify the samples collected in different days and identify metabolic differences during the treatment. In OPLS-DA separation between two classes was forced in the first component (the “predictive” component) and further components, orthogonal to the first, only expressed intra-class variability. The classification power of models is expressed by the correlation coefficient R^2^Y. 7-fold cross-validation was performed to optimize model parameters, whereas permutation test on the responses (1000 random permutations) was used to highlight the presence of over-fitting according to good practice for model building.

Using the Chenomx NMR Suite Profiler tool, main metabolites were quantified. Urinary concentrations were then adjusted to the 24 h urinary output to obtain absolute metabolite values (μg/24h). Metabolomes were compared across study groups (i.e. those receiving QD, BID, TID, and control) using a mixed-effects, restricted maximal likelihood estimation regression, with repeated measures occurring over days using Stata version 16.1 (Statacorp LLC, College Station, TX). Measures were repeated at the level of the individual rat. Spearman correlation was performed for injury biomarkers and metabolome. All tests were two-tailed, with an a priori level of statistical significance set at an alpha of 0.05.

## RESULTS

^1^H NMR experiments were performed on all the samples (i.e. n=77). Main low molecular weight metabolites were identified, including amino acids, organic acids, osmolytes, and sugars. In order to extract the latent information contained in the NMR spectra and identify the metabolomic profiles associated to the different phases of the experiment, spectral data were submitted to multivariate statistical analysis.

An exploratory unsupervised PCA was first performed on urinary spectral data to visualize trends and clustering among similar samples and identify possible outliers. The first three PCs accounted for 71.0% of the total variance. In the PCA score scatter plot (Fig. 1A), the urine samples were clearly separated into three groups according to the time of collection, namely pre-surgery (day 0), dosing day 1, and dosing day 3 samples, indicating that they were characterized by different metabolomic profiles, independent of the PB fractionation scheme. Day 0 samples lie in the lower left quadrant and were extremely homogeneous forming a tight cluster, indicative of a very similar profile. This is consistent with the uniform animal model chosen and that all animals had not entered the treatment protocol yet. Post treatment sample clusters were less homogeneous indicating a higher interindividual variability in response to the PB dose, with day 1 and day 3 clearly separated. Moreover, the two post 1^st^ dose samples lying in the multivariate space near to the day 0 ones correspond to the urinary samples of two sham animals collected after a saline dose. This contiguity indicates similar metabolomic profiles, even though sham animals are clearly distinguishable from the others due to the surgical procedure. The analysis of the loading plot reported in Fig. 1B indicates which variables (metabolites) were responsible for the sample distribution in the score plot: day 0 samples were characterized by higher amounts of citrate, 2-oxoglutarate, and hippurate, while day 1 samples show higher taurine, and day 3 samples higher lactate, acetate and creatine.

**Fig 1.**
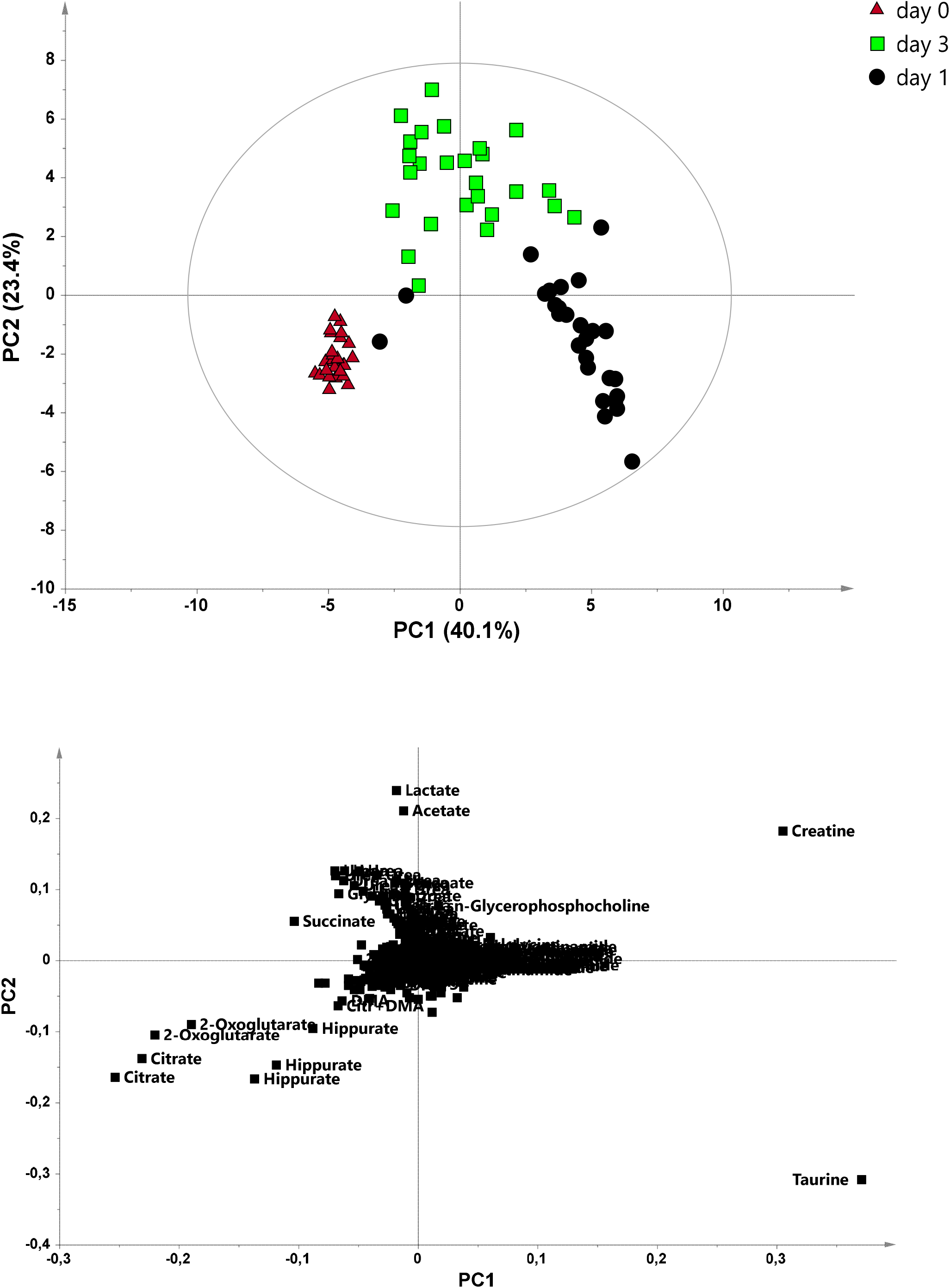
PCA model of all the analyzed urine samples (A=3, Par, R^2^=0.71, Q2=0.60). a) Score plot of the samples colored by day; b) loading plot of the variables (metabolites).

To classify the samples belonging to the different groups and to identify the corresponding discriminant metabolites, a supervised OPLS-DA analysis was conducted, and different OPLS-DA models were built comparing two groups at a time. Fig. 2 shows the OPLS-DA model of day 0 vs. day 1 samples. The predictive component (tp, reported in the x-axis) accounts for the difference between the two studied groups, while the orthogonal component (to1, y-axis) accounts for the intragroup variability. The two groups were well separated, further indicating a different metabolomic profile. Moreover, day 0 samples showed a very low intragroup variability, while day 1 samples were distributed along the orthogonal component, the distribution being related to the relative amount of taurine, as indicated by the sample coloring. Interestingly, taurine distribution in day 1 samples was also related to the PB fractionation scheme; QD and most of the BID samples were distributed in the lower part of the multivariate space and were characterized by relatively lower taurine levels, compared to TID samples which were distributed in the upper part and showed higher taurine levels (with the exception of one BID group animal). Differences between metabolites on day 1 and day 0 samples are reported in Table 1 and were calculated for each of the dosing schemes. Raw values of changes per individual are visualized in Supplemental figure 1.

**Fig 2.**
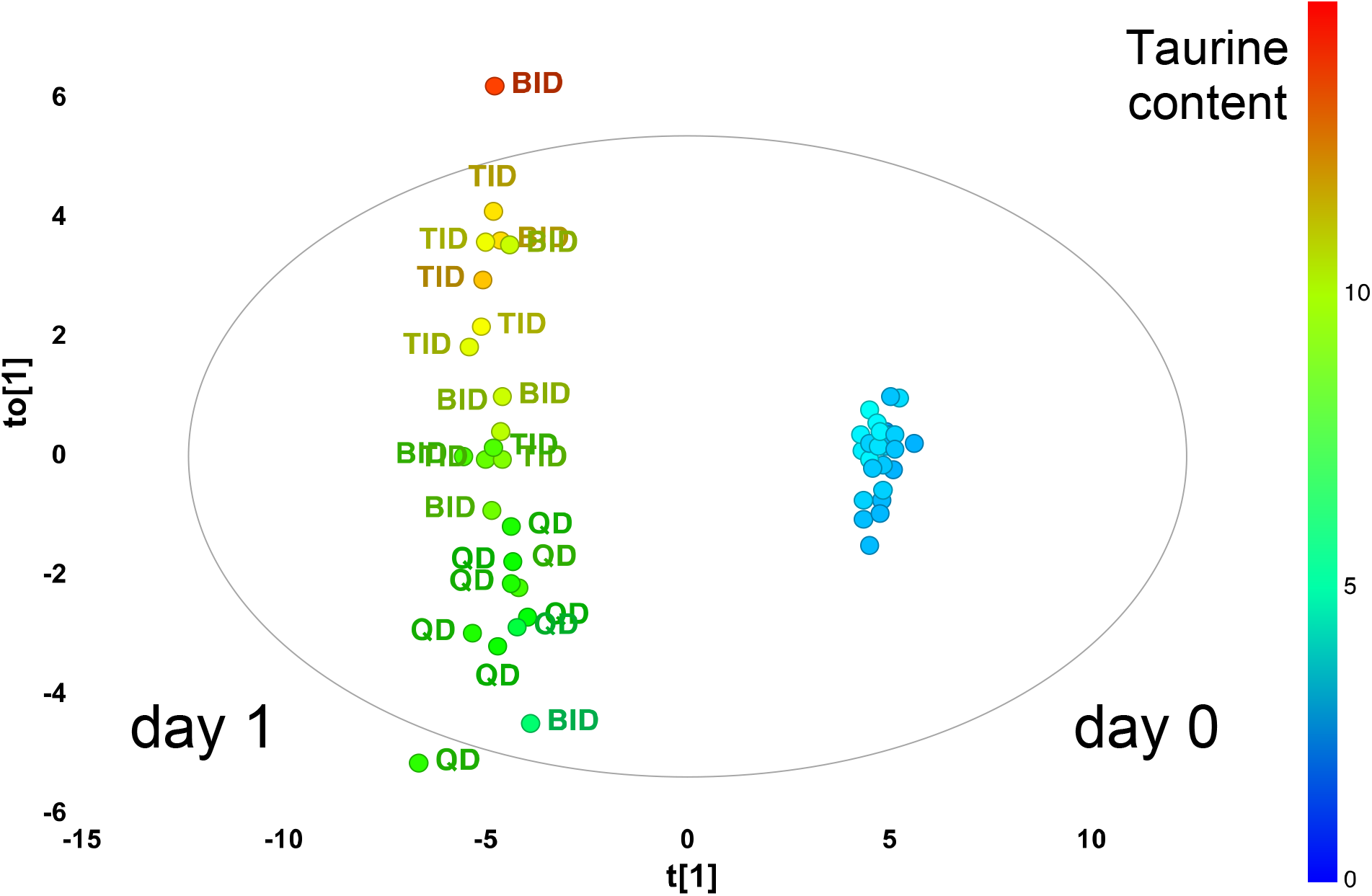
OPLS-DA score plot of urine samples at day 0 vs day 1. A=1+1, Par, R^2^Y=0.98, Q2Y=0.98. Samples are colored according to the taurine content and labelled according to the dosing scheme.

**Table 1.** Statistical differences between metabolites (μg/24h) on day 1 vs day 0 between treatment groups QD, BID, and TID

|  | Metabolite | QD | BID | TID |
| --- | --- | --- | --- | --- |
| **Higher at day 1** | Taurine | <0.001 | <0.001 | <0.001 |
|  | Creatine | <0.001 | <0.001 | <0.001 |
| **Lower at day 1** | β-alanine | <0.001 | <0.001 | 0.001 |
|  | Betaine | 0.008 | <0.001 | <0.001 |
|  | Choline | 0.004 | <0.001 | 0.007 |
|  | Citrate | <0.001 | <0.001 | <0.001 |
|  | Creatinine | 0.115 | <0.001 | 0.483 |
|  | Dimethylsulfone | <0.001 | <0.001 | <0.001 |
|  | Glutamine | 0.003 | <0.001 | 0.012 |
|  | Hippurate | <0.001 | <0.001 | <0.001 |
|  | 2-Oxoglutarate | <0.001 | <0.001 | <0.001 |
|  | Pyruvate | <0.001 | <0.001 | <0.001 |
|  | Succinate | <0.001 | <0.001 | <0.001 |
|  | Sucrose | <0.001 | <0.001 | <0.001 |
|  | TMAO | <0.001 | <0.001 | <0.001 |
|  | DMA | <0.001 | <0.001 | NS |
|  | Methionine | 0.016 | <0.001 | NS |
|  | Glycine | NS | 0.002 | NS |
|  | Leucine | NS | <0.001 | NS |
|  | Valine | NS | 0.010 | NS |
|  | Acetoacetate | NS | NS | 0.050 |
|  | Isoleucine | NS | NS | 0.022 |

Fig. 3 shows the OPLS-DA model of day 0 vs. day 3 samples. The two groups were well separated, and as previously, day 3 samples were stratified along the orthogonal component; the distribution in this case was related to the relative amount of creatine, as indicated by the sample coloring, whereas no correlation was observed with the PB dosing scheme. Statistical differences between metabolites on day 3 and day 0 samples are reported in Table 2 and were calculated for each of the dosing schemes. Fig. 4 shows the OPLS-DA model of day 1 vs. day 3 samples. The two groups were well separated, and each group shows a high intragroup variability.

**Fig 3.**
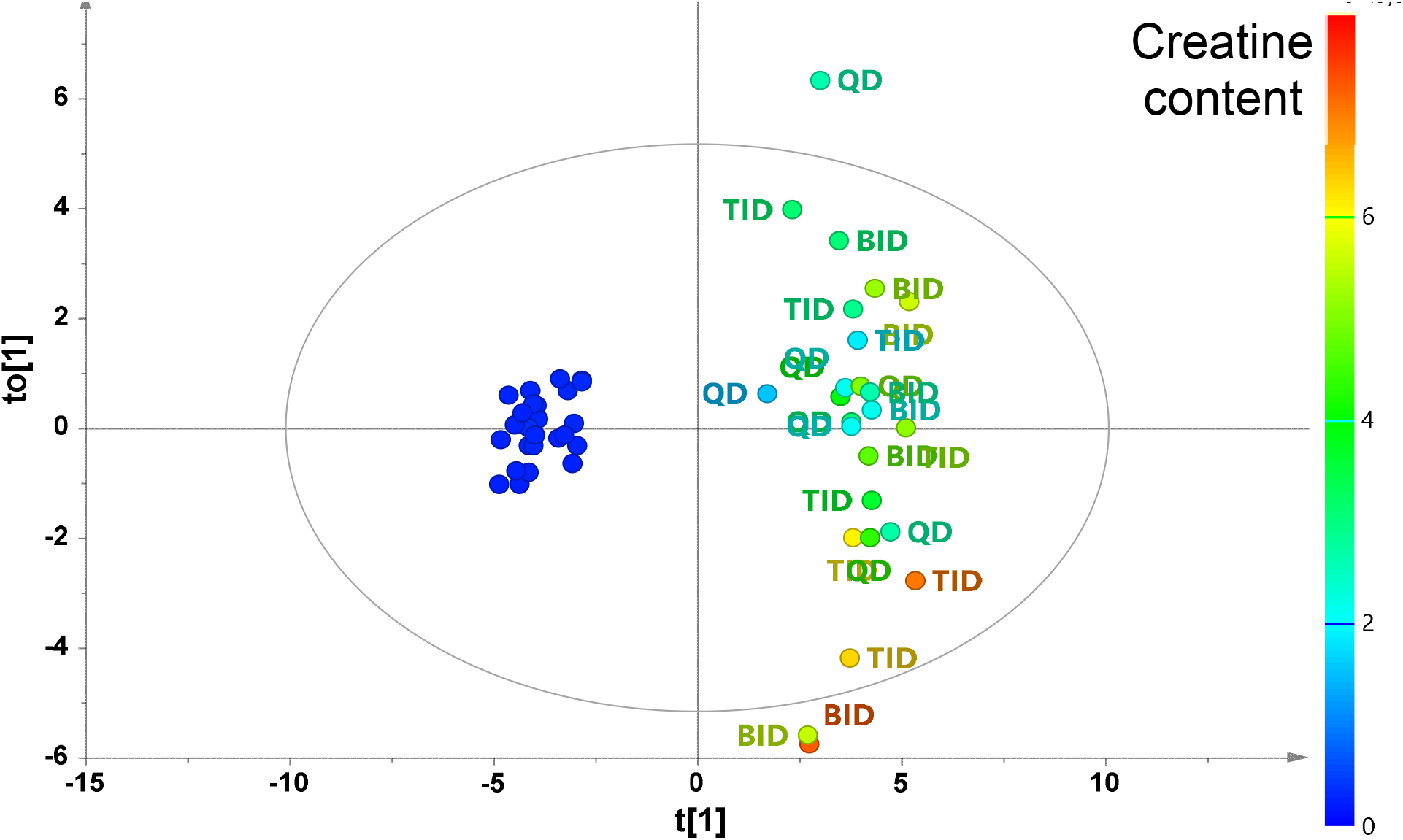
OPLS-DA score plot of urine samples at day 3 vs day 0 (A=1+1, Par, R^2^Y=0.96, Q^2^Y=0.95). Samples are colored according to the creatine content and labelled according to the dosing scheme.

**Table 2.**
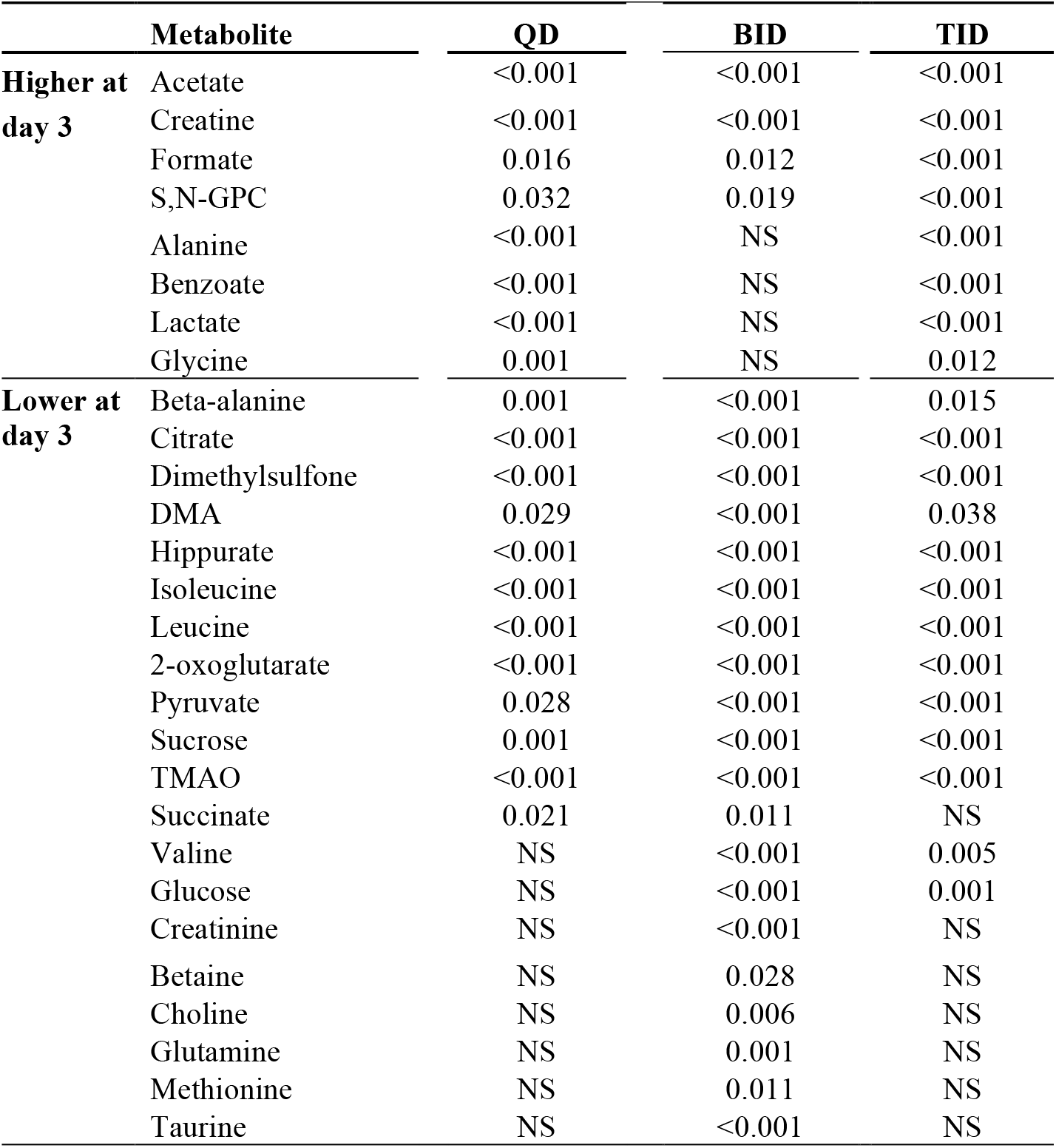
Statistical differences between metabolites (μg/24h) on day 3 vs day 0 between treatment groups QD, BID, and TID

**Fig 4.**
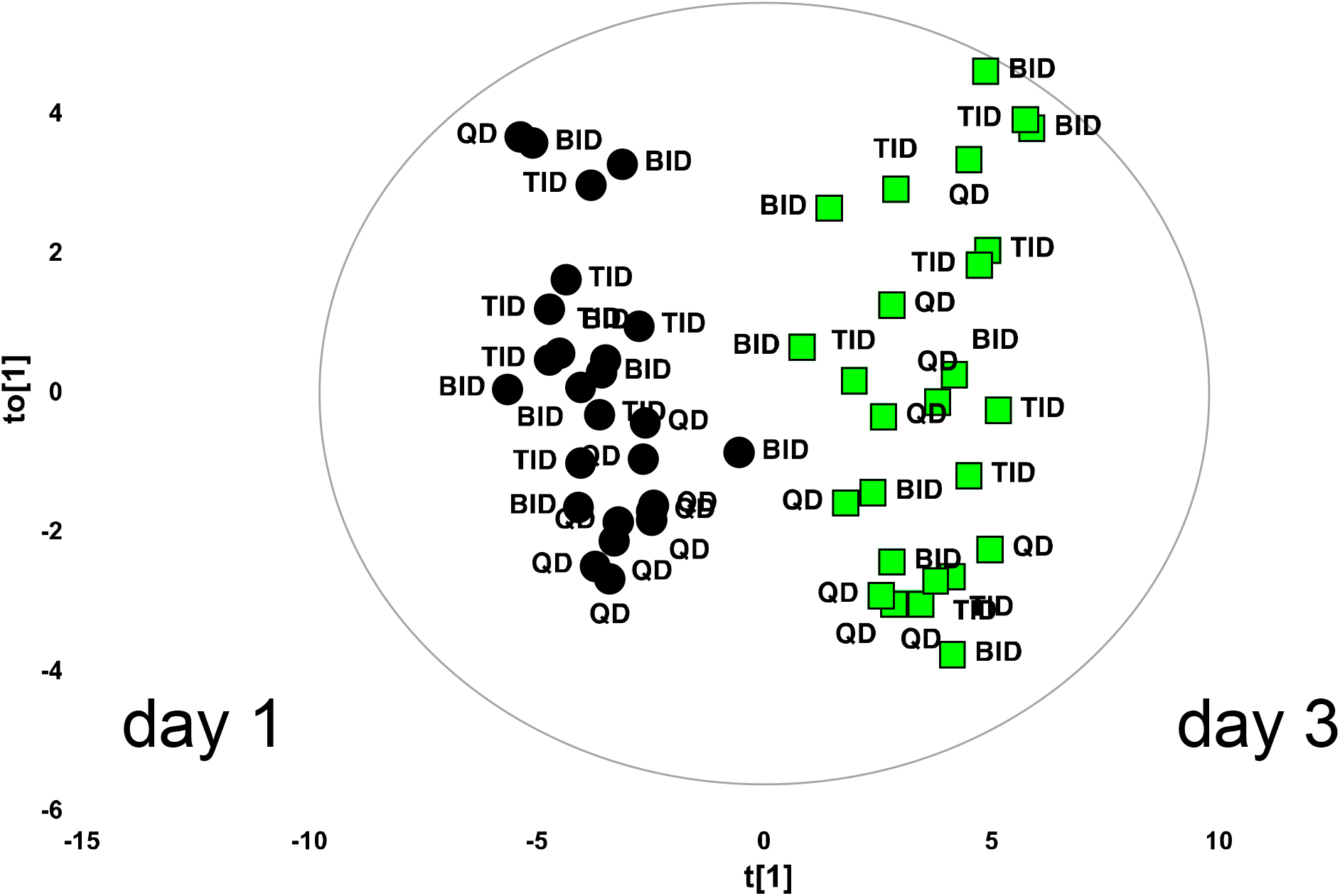
OPLS-DA score plot of urine samples at day 3 vs day 1. A=1+1, Par, R^2^Y=0.84, Q^2^Y=0.80.

Metabolites characterizing day 3 urinary metabolome profile to day 1 are reported in Table 3. When comparing day 1 to day 3, the following metabolites were higher: lactate, acetate, succinate, glycine, citrate, 2-oxoglutarate, benzoate, choline, SN-GPC, alanine, formate, and sucrose. Conversely, taurine, hippurate, isoleucine, and leucine were lower (Supplementary Figure 1). When the analysis was performed using QD dosing scheme as a reference, the TID group showed a more polymorphic response on day 1, compared with the BID group (see Table 3). Taurine was the only metabolite significantly increased in both BID and TID compared to QD group. Few other metabolites contribute to the differentiation of dosing schemes over time. The metabolite differences among the different dosing groups at day 3 are reported in Table 3. Taurine was the most discriminant metabolite as a biological marker related to both PB administration and dosing scheme on day 1; notably a correlation between urinary KIM-1 concentration and taurine measured on the same samples was observed on day 1 (Spearman’s rho =0.4052, P=0.036, Fig. 5a). If focused on day 3, few other metabolites seem to be more relevant for the metabolomic profile characterization (Fig. 5c-f). Finally, when looking at early predictors for histopathologic kidney injury, taurine showed promise on day 1 for correlating with increasing histopathology scores (Spearman’s rho = 0.4167, P=0.038) whereas KIM-1 on day one and day 3 did not reach significance (Spearman’s rho = 0.3248, P=0.11 and Spearman’s rho = 0.3739, P=0.066, respectively and shown in Figure S2a and S2b).

**Table 3.**
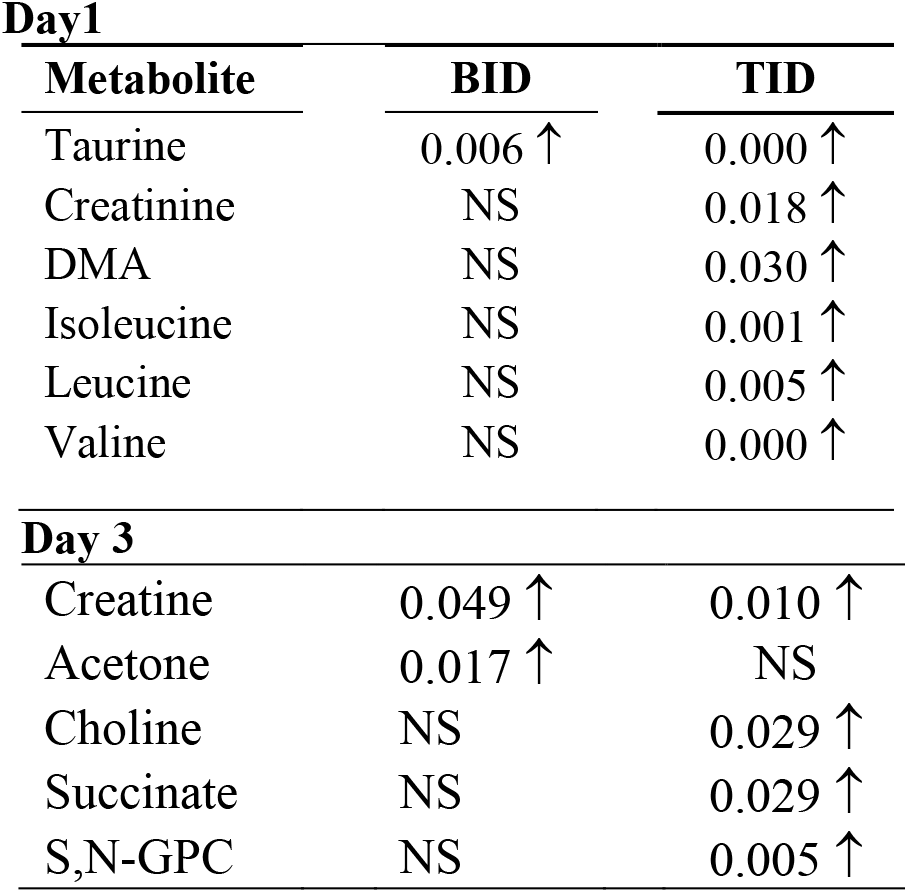
Statistical differences between metabolites (μg/24h) on day 1 and day 3 between treatment groups (BID and TID vs QD).

**Day1**
| Metabolite | BID | TID |
| --- | --- | --- |
| Taurine | 0.006 ↑ | 0.000 ↑ |
| Creatinine | NS | 0.018 ↑ |
| DMA | NS | 0.030 ↑ |
| Isoleucine | NS | 0.001 ↑ |
| Leucine | NS | 0.005 ↑ |
| Valine | NS | 0.000 ↑ |
| **Day 3** |  |  |
| Creatine | 0.049 ↑ | 0.010 ↑ |
| Acetone | 0.017 ↑ | NS |
| Choline | NS | 0.029 ↑ |
| Succinate | NS | 0.029 ↑ |
| S,N-GPC | NS | 0.005 ↑ |

**Fig 5.**
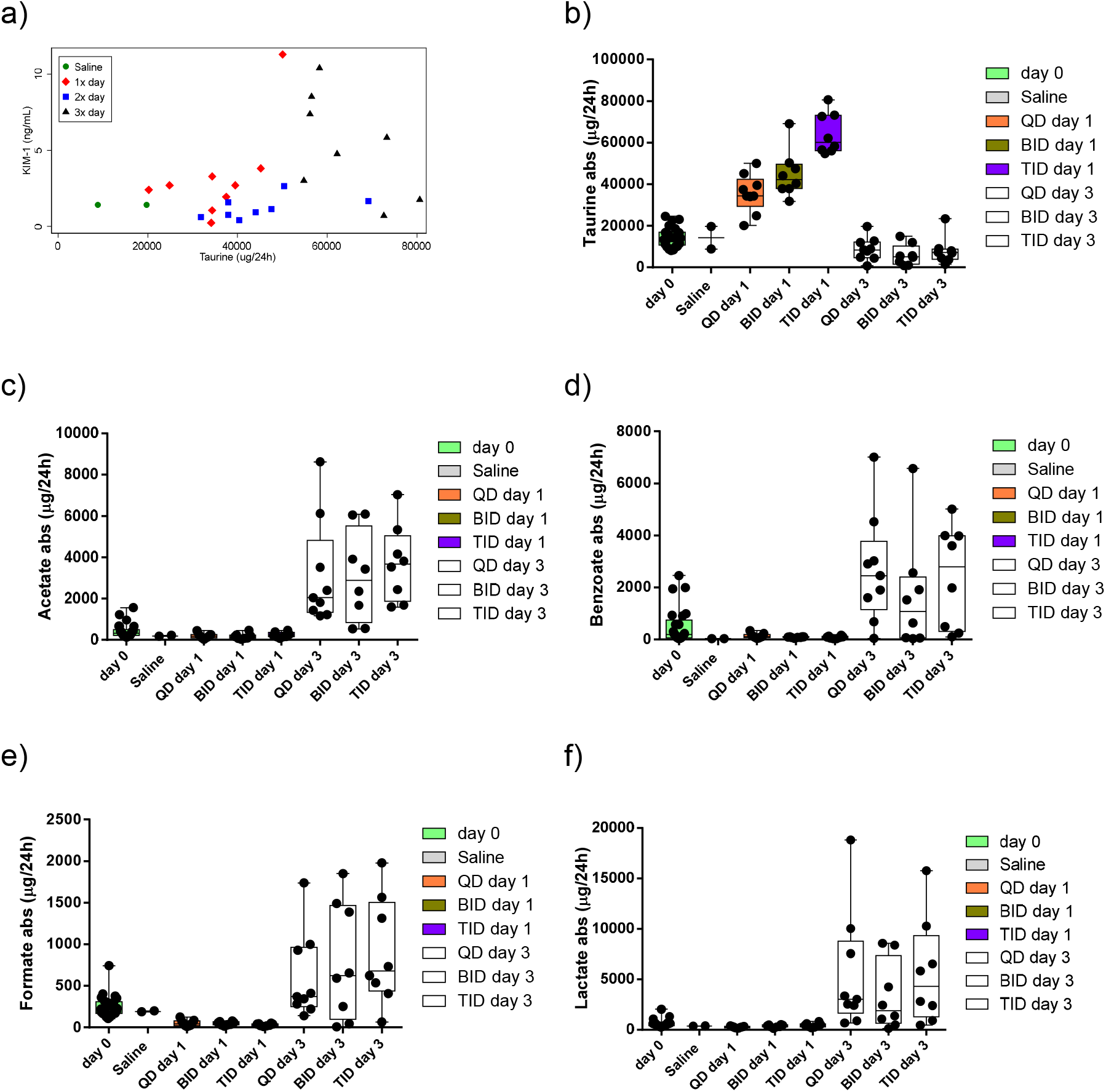
a) Day 1, KIM-1 vs. Taurine correlation (P=0.04), b) taurine box plot at different collection timepoints and PB dosing schemes, and c-f) box plots of metabolites characterizing urinary profile at day 3.

## DISCUSSION

In this longitudinal metabolomic profile analysis of three PB dosing regimens administered for 3 days in a healthy rat model, we found increased amounts of urinary taurine on day 1 after dosing with PB, regardless of fractionation schedule. Taurine increased with KIM-1 on day 1 after dosing and then decreased back to baseline whereas KIM-1 increased throughout the treatment.^14^ KIM-1 is a known renal injury biomarker that correlates with proximal tubule damage from xenobiotics as a function of being involved in the signaling of phagocytotic clearance of apoptotic debris.^25–27^ Thus, taurine may represent a physiological response to the nephrotoxic effects of PB whereas KIM-1 serves as a marker of damage (and host immune repair). Elucidation of taurine expression in response to kidney injury may help clinicians identify polymyxin-associated nephrotoxicity earlier.

In our experiment, the urinary metabolome profiles at baseline (day 0) clustered together, being characterized by higher levels of citrate, 2-oxoglutarate, and hippurate when compared with urinary profiles of the following days (day 1 and day 3). We interpret the comparatively higher levels of these metabolites at baseline as a ‘relative’ behavior, being the results both of longitudinal cellular consumption and active/passive release/reuptake into the urine. It was clear that subcutaneous administration of PB to rats, independently from concentration and time effects, caused a clear modification of metabolomic profiles in treated animals. The overview of the PCA analysis, which takes into account all the profiles in the three days under analysis, demonstrates the differences exhibited by the urinary profiles following the first dose of polymyxin. The baseline profiles are clustered together, with clear separation from the day 1 and day 3 profiles. When a day-by-day comparison is performed by a supervised approach (OPLS-DA), a set of metabolites was found to contribute to this separation. After the first dose – in a temporal window of nearly 24 hours – taurine drives the clear differentiation from both baseline and post-3^rd^ dose urinary metabolomic profiles.

A previous study by Azad et al utilized a metabolomic approach in their investigation of PB-induced toxicity in rat kidney proximal tubule cells (NRK-52E).^28^ Both concentration and time-dependent transitions of mitochondrial morphology were observed in NRK-52E cells following PB treatment (1 and 2 mM at intervals up to 24 h). Disruption of the taurine-hypotaurine pathway in polymyxin-treated kidney cells was also found to be associated with polymyxin-induced toxicity in the form of impaired cellular ROS scavenging. This metabolite, and its metabolic pathway, was identified as one of the main metabolic perturbations related to PB-induced tubular kidney cell (HK-2 and NRK-52E) damage.^29^ The underlying mechanism of damage was assumed to be, among the aforementioned, oxidative and related to a loss of cellular capacity to scavenging ROS. The sharp surge of urinary taurine in all of the treated animals in our study is limited to the day 1 profile, with a return to baseline by the day 3 profile; indicative of a prompt cellular response to the accumulation of PB inside the proximal tubule cells after its administration. Taurine efflux from proximal tubule cells and its subsequent excretion in urine, is on average much slower than its uptake (due to its higher K_m_)^30^, being dependent on its inner cellular concentration, on osmotic forces (acting as an important urinary osmolyte), and on a carrier-facilitated process, as for its active uphill transport. Its rapid release in urine may be explained by active excretion from proximal tubule cells into the lumen and eventually into the urine, or by a passive discharge due to proximal tubule cell necrosis or to apoptotic death. The latter mechanism would imply the presence in urine of a conspicuous amount of all the other metabolite actively concentrated into the proximal tubule cells, but this was not the case. It is possible that taurine, which has antioxidant, cytoprotective, and osmolyte activity, counteracts the injurious effects of PB on proximal tubule cells.^30^ Taurine may function as a first response in this regard, and its stores may be quickly depleted (in the setting of continued/prolonged injury), causing urinary taurine levels to return to baseline. In relation to our findings, a previous investigation of exogenous taurine for amelioration of colistin-induced nephrotoxicity found that taurine supplementation decreased both oxidative stress and mitochondrial dysfunction in renal tissue.^31^

When comparing the metabolomics profiles of post-1^st^ dose and post-3^rd^ dose, the urinary profile after the third dose is characterized by the presence of several TCA cycle intermediates (namely, succinate, citrate, and 2-oxoglutarate) of anaerobic glycolysis (lactate) and membrane damage (choline). The higher amounts of baseline citrate, 2-oxoglutarate (both intermediates of Krebs/TCA cycle), and hippurate may be interpreted as existing at a baseline level, with subsequent decline on days 1 and 3 reflective of their depletion under cellular stress. Holmes et al. investigated the urinary effects of several kidney damaging substances by a metabolomic approach and found similar high levels of these metabolites in baseline urinary metabolomic profiles, stating that their subsequent decreases (mainly by consumption and lower production) were the shared consequence of several damaging mechanisms in the kidney.^32^

Our work provides an early characterization of the urinary metabolomics related to polymyxin-associated nephrotoxicity. The results are confirmatory with respect to the identification of urinary metabolites that are expressed early on in renal toxicity and novel in regards to characterization of the metabolome after PB dosing. In particular, correlation of elevated urinary taurine levels to renal injury biomarkers (KIM-1) and characterization of the urinary metabolomic profile in the presence of nephrotoxins may have important clinical implications. Although further study is still needed, these early results may ultimately aid in the earlier detection of polymyxin-associated nephrotoxicity.

## CONCLUSION

In summary, we demonstrated that PB causes increased amounts of urinary taurine on day 1 which then normalizes to baseline concentrations. The initial taurine response is correlated with KIM-1, a well-known proximal tubule kidney injury biomarker, indicating that taurine could be involved as a physiologic protective response to the insult by PB. Taurine may provide one of the earlier signals of acute kidney damage caused by PB. Our metabolomic approach to classifying ‘polymyxin nephrotoxicity’ has revealed several mechanistic changes that correspond to kidney injury and may help elucidate ameliorating agents for renal tubular cell injury. Future work should consider taurine as a potential ameliorating agent given pre-emptively for patients receiving PB.

## Supplementary figures

**Figure S1.**
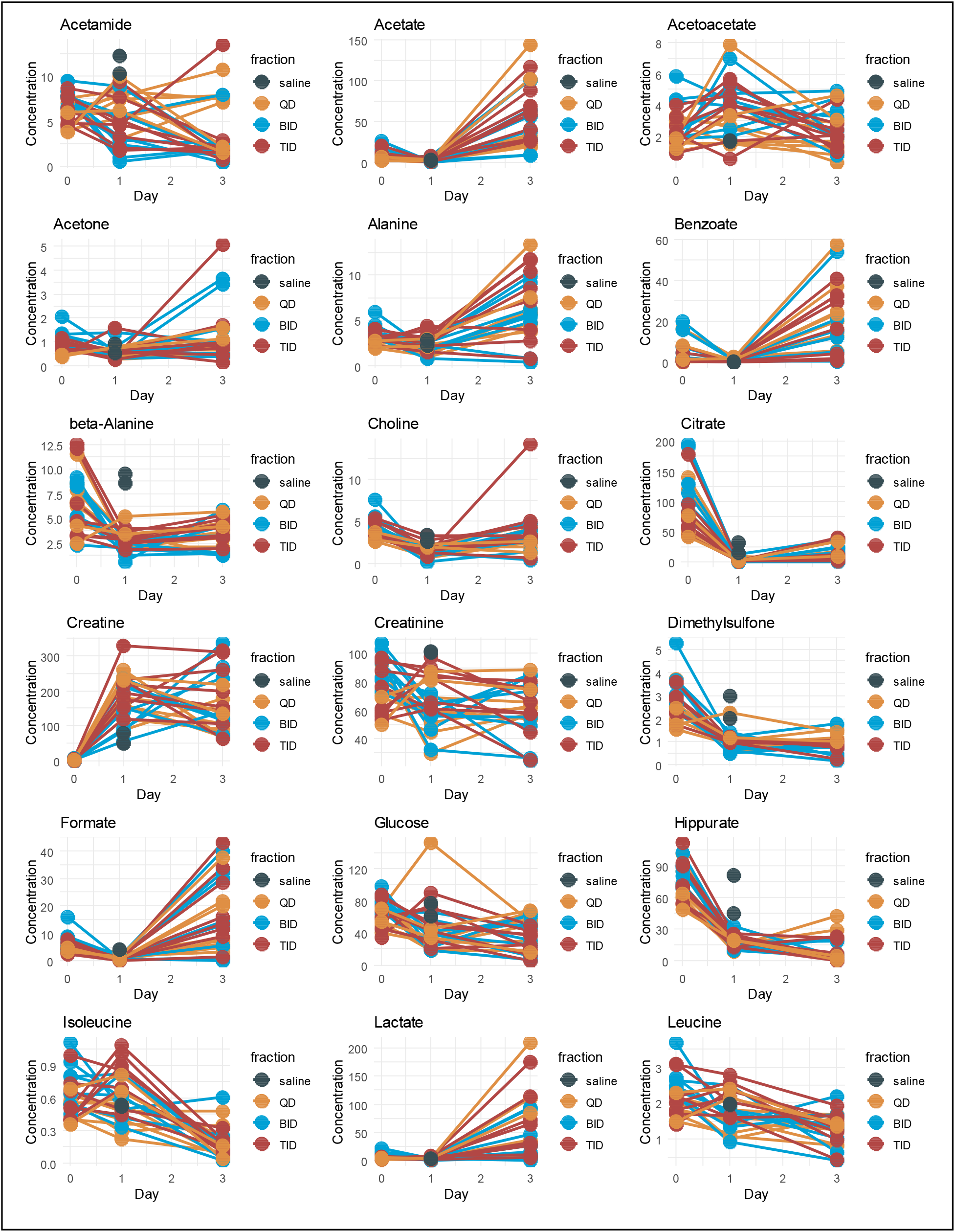
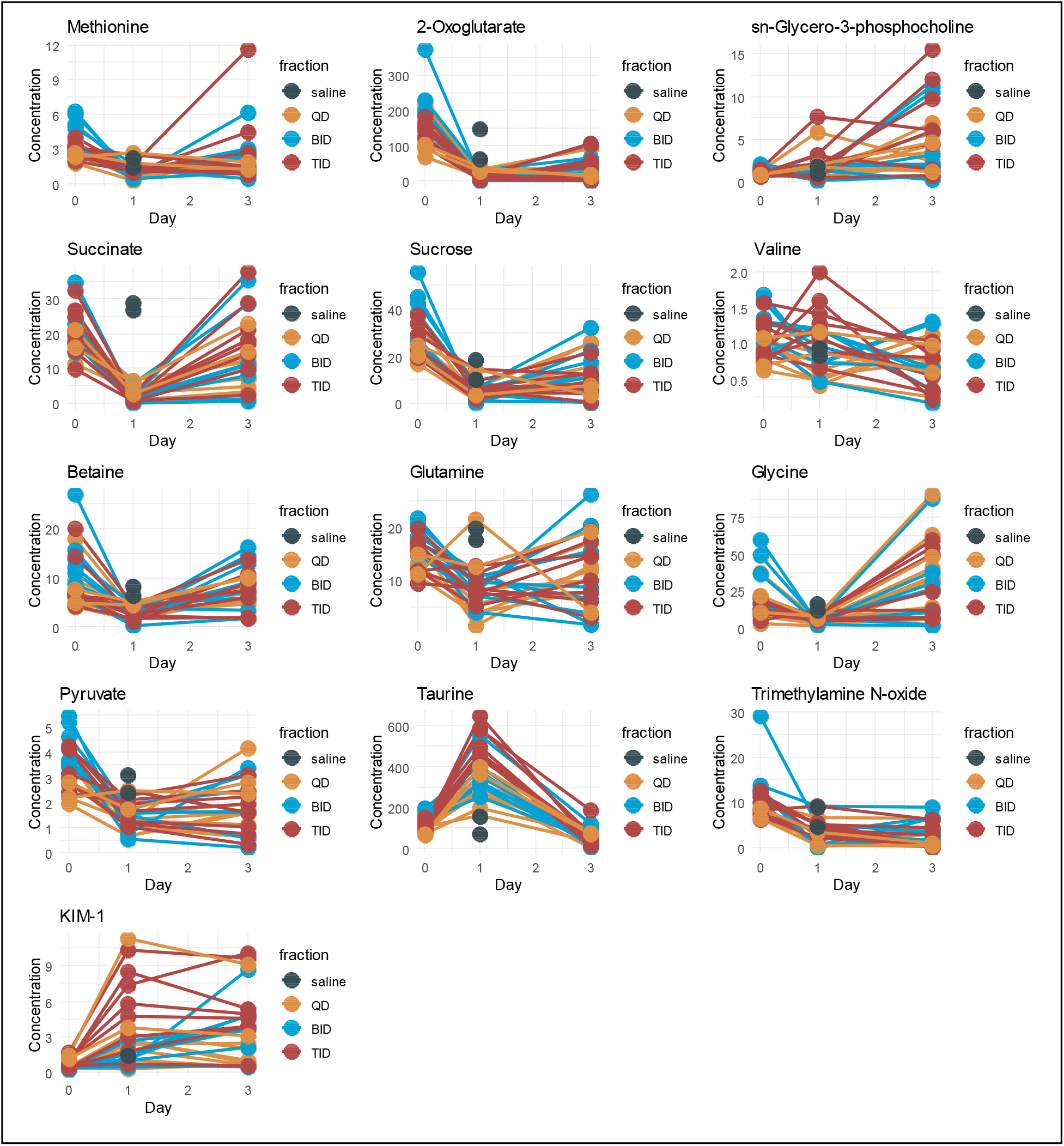

**Figure S2.**
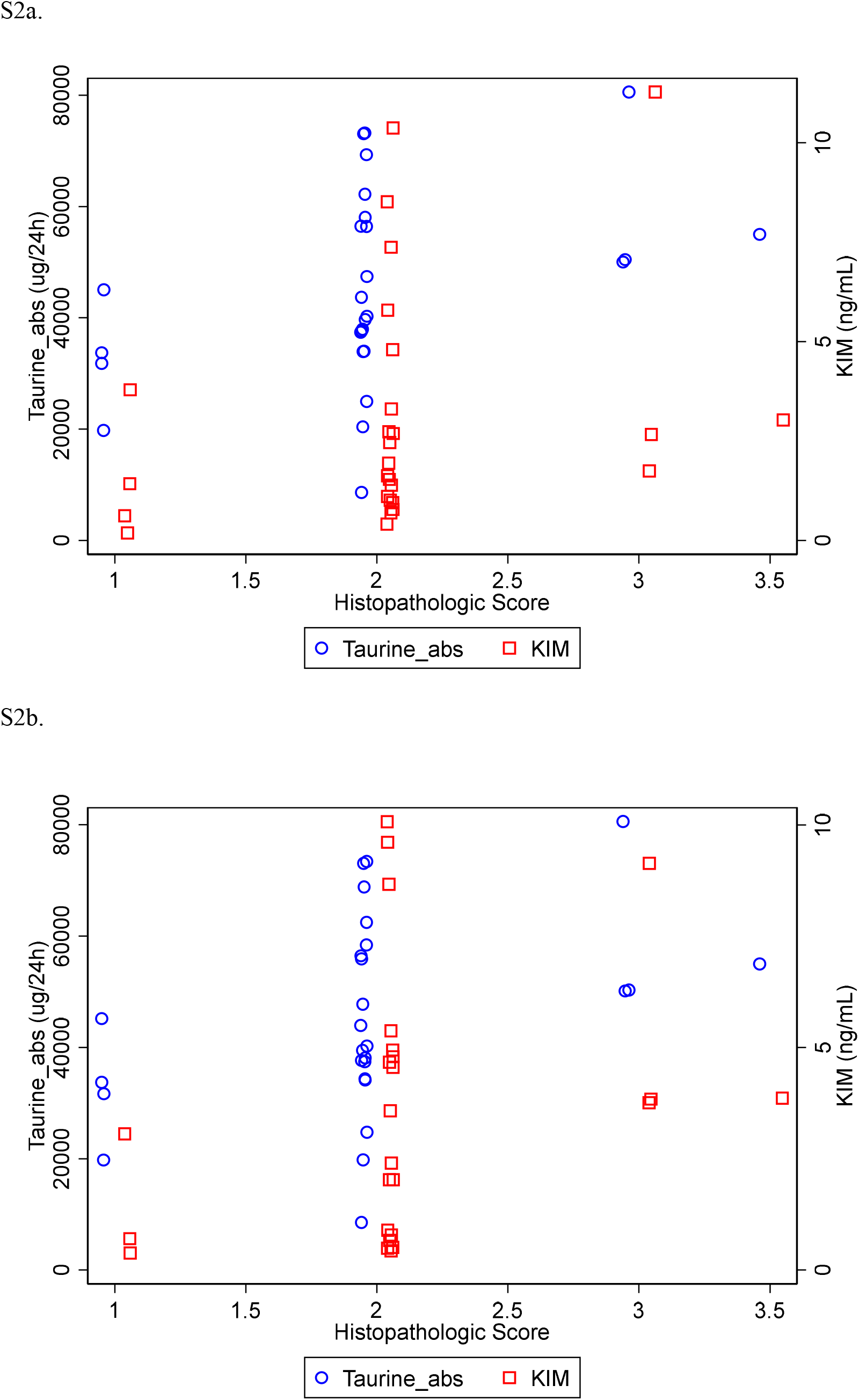
Urinary KIM-1 (day 1) and Taurine (day 1) versus final histopathological score (S2a) and Urinary KIM-1 (day 3) and Taurine (day 1) versus final histopathological score (S2b). Mild jitter applied to visualize identical results. Taurine correlated with increasing histopathology scores on day 1 (Spearman’s rho = 0.4167, P=0.038) whereas KIM-1 on day 1 and day 3 did not reach significance (Spearman’s rho = 0.3248, P=0.11 and Spearman’s rho = 0.3739, P=0.066). S2a.

## REFERENCES

1. WHO Team. 2020 antibacterial agents in clinical and preclinical development: an overview and analysis, 2020.

2. Klein EY, Van Boeckel TP, Martinez EM et al. Global increase and geographic convergence in antibiotic consumption between 2000 and 2015. Proceedings of the National Academy of Sciences 2018; 115: E3463–E70.

3. Yun B, Azad MAK, Wang J et al. Imaging the distribution of polymyxins in the kidney. The Journal of antimicrobial chemotherapy 2015; 70: 827–9.

4. Azad MAK, Roberts KD, Yu HH et al. Significant Accumulation of Polymyxin in Single Renal Tubular Cells: A Medicinal Chemistry and Triple Correlative Microscopy Approach. Analytical Chemistry 2015; 87: 1590–5.

5. Lu X, Chan T, Xu C et al. Human oligopeptide transporter 2 (PEPT2) mediates cellular uptake of polymyxins. The Journal of antimicrobial chemotherapy 2016; 71: 403–12.

6. Manchandani P, Zhou J, Babic JT et al. Role of Renal Drug Exposure in Polymyxin B-Induced Nephrotoxicity. Antimicrobial agents and chemotherapy 2017; 61: e02391–16.

7. Sivanesan SS, Azad MAK, Schneider EK et al. Gelofusine Ameliorates Colistin-Induced Nephrotoxicity. Antimicrobial agents and chemotherapy 2017; 61: e00985–17.

8. Azad MAK, Nation RL, Velkov T et al. Mechanisms of Polymyxin-Induced Nephrotoxicity. Advances in experimental medicine and biology 2019; 1145: 305–19.

9. Nation RL, Rigatto MHP, Falci DR et al. Polymyxin Acute Kidney Injury: Dosing and Other Strategies to Reduce Toxicity. Antibiotics (Basel, Switzerland) 2019; 8: 24.

10. Falagas ME, Kasiakou SK. Colistin: the revival of polymyxins for the management of multidrug-resistant gram-negative bacterial infections. Clin Infect Dis 2005; 40: 1333–41.

11. Avedissian SN, Liu J, Rhodes NJ et al. A Review of the Clinical Pharmacokinetics of Polymyxin B. Antibiotics (Basel, Switzerland) 2019; 8: 31.

12. Abdelraouf K, Braggs KH, Yin T et al. Characterization of polymyxin B-induced nephrotoxicity: implications for dosing regimen design. Antimicrob Agents Chemother 2012; 56: 4625–9.

13. Okoduwa A, Ahmed N, Guo Y et al. Nephrotoxicity Associated with Intravenous Polymyxin B Once-versus Twice-Daily Dosing Regimen. Antimicrob Agents Chemother 2018; 62.

14. Liu J, Pais GM, Avedissian SN et al. Evaluation of Dose-Fractionated Polymyxin B on Acute Kidney Injury Using a Translational *In Vivo* Rat Model. Antimicrobial Agents and Chemotherapy 2020; 64: e02300–19.

15. Sun W, Hu B, Zhang X et al. Effect of Different Dosage Frequency of Polymyxin B on Rat Nephrotoxicity. Drug design, development and therapy 2021; 15: 611–6.

16. Ozyilmaz E, Ebinc FA, Derici U et al. Could nephrotoxicity due to colistin be ameliorated with the use of N-acetylcysteine? Intensive Care Med 2011; 37: 141–6.

17. Yousef JM, Chen G, Hill PA et al. Ascorbic acid protects against the nephrotoxicity and apoptosis caused by colistin and affects its pharmacokinetics. J Antimicrob Chemother 2012; 67: 452–9.

18. Yousef JM, Chen G, Hill PA et al. Melatonin Attenuates Colistin-Induced Nephrotoxicity in Rats. Antimicrobial Agents and Chemotherapy 2011; 55: 4044–9.

19. Azad MAK, Sivanesan S, Wang J et al. Methionine Ameliorates Polymyxin-Induced Nephrotoxicity by Attenuating Cellular Oxidative Stress. Antimicrob Agents Chemother 2018; 62.

20. Waikar SS, Bonventre JV. Creatinine kinetics and the definition of acute kidney injury. J Am Soc Nephrol 2009; 20: 672–9.

21. Ronco C, Rosner MH. Acute kidney injury and residual renal function. Crit Care 2012; 16: 144.

22. Lassnigg A, Schmidlin D, Mouhieddine M et al. Minimal changes of serum creatinine predict prognosis in patients after cardiothoracic surgery: a prospective cohort study. J Am Soc Nephrol 2004; 15: 1597–605.

23. Sistare FD, Dieterle F, Troth S et al. Towards consensus practices to qualify safety biomarkers for use in early drug development. Nature Biotechnology 2010; 28: 446–54.

24. Wishart DS, Feunang YD, Marcu A et al. HMDB 4.0: the human metabolome database for 2018. Nucleic Acids Research 2017; 46: D608–D17.

25. Vaidya VS, Ozer JS, Dieterle F et al. Kidney injury molecule-1 outperforms traditional biomarkers of kidney injury in preclinical biomarker qualification studies. Nature Biotechnology 2010; 28: 478–85.

26. Bonventre JV. Kidney injury molecule-1 (KIM-1): a urinary biomarker and much more. Nephrology Dialysis Transplantation 2009; 24: 3265–8.

27. Ichimura T, Asseldonk EJPv, Humphreys BD et al. Kidney injury molecule–1 is a phosphatidylserine receptor that confers a phagocytic phenotype on epithelial cells. The Journal of Clinical Investigation 2008; 118: 1657–68.

28. Azad MAK, Akter J, Rogers KL et al. Major Pathways of Polymyxin-Induced Apoptosis in Rat Kidney Proximal Tubular Cells. Antimicrobial Agents and Chemotherapy 2015; 59: 2136–43.

29. Nang SC, Azad MAK, Velkov T et al. Rescuing the Last-Line Polymyxins: Achievements and Challenges. Pharmacological Reviews 2021; 73: 679–728.

30. Chesney RW, Han X, Patters AB. Taurine and the renal system. Journal of biomedical science 2010; 17 Suppl 1: S4–S.

31. Heidari R, Behnamrad S, Khodami Z et al. The nephroprotective properties of taurine in colistin-treated mice is mediated through the regulation of mitochondrial function and mitigation of oxidative stress. Biomed Pharmacother 2019; 109: 103–11.

32. Holmes E, Nicholls AW, Lindon JC et al. Development of a model for classification of toxin-induced lesions using 1H NMR spectroscopy of urine combined with pattern recognition. NMR in Biomedicine 1998; 11: 235–44.

